# Mechanism of biomimetic virus-like nanoparticles in triggering immune response elucidated by proteomics

**DOI:** 10.1101/2025.02.27.640536

**Authors:** Jiajia Wang, Ahmed Montaser, Minna Sivonen, Henri Leinonen, Jussi Hepojoki, Leyuan Ma, Vesa-Pekka Lehto, Wujun Xu

## Abstract

The use of biomimetic nanoparticles (NPs) with virus-like morphology have recently attracted research interest as novel delivery platforms and immune adjuvants. However, the exact interactions between the nanoparticles and immune cells as well as the mechanism involved are not known in detail. This motivated us to develop virus-like mesoporous silica nanoparticles (VLP) to characterize their physicochemical properties, and to determine the immune pathways induced by the particles in mouse macrophages. The results showed inclusion of spikes mimicking virion structures on the surface increased cellular uptake and enhanced immune response as compared to spherical NPs. Proteomic analysis revealed that the RIG-I-like receptor signaling pathway, Chemokine signaling pathway, MAPK signaling pathway, NF-κB signaling pathway, Toll-like receptor signaling pathway, B cell receptor signaling pathway and Th1 and Th2 cell differentiation pathways were involved in regulating the immune response when macrophages interacted with VLP. When the spikes increased from 5 to 30 nm, the expression levels of immune-related proteins including TRAF6 and PIAS4 proteins enhanced. This study revealed the interaction pathways and key proteins in the activation of immune response with VLP, which may provide insights to develop novel immunotherapy for enhanced efficacy.

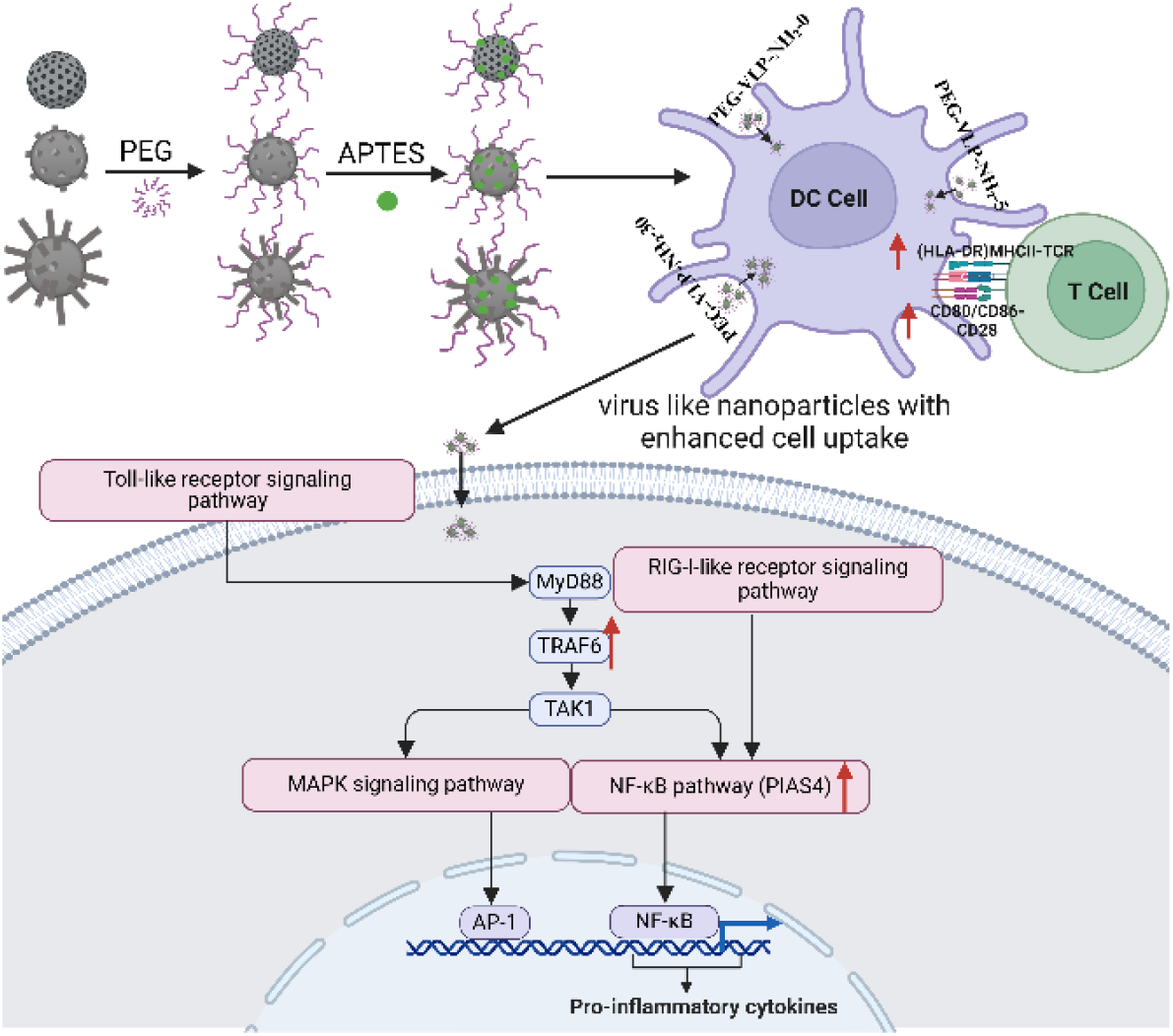

## Introduction

Mesoporous silica nanoparticles (MSNs) have gained significant attention in cancer therapy due to their unique properties, such as uniform mesoporous structure, large surface area, good biocompatibility, high loading capacity, and easy to achieve functionalization^1,2^. For cancer immunotherapy, MSNs have been explored as immune adjuvants to stimulate both innate and adaptive immune responses^3^. In addition, due to their high surface area, they are good nanocarriers for delivering various therapeutic antigens used in cancer vaccines^4^. For example, MSNs enhanced the intracellular delivery of R848, a Toll-like receptor agonist into the lysosomes of antigen-presenting cells (APCs), thereby effectively activating dendritic cells (DCs) and antigen-specific T cell responses^5^. Furthermore, MSNs have outstanding performance in the co-delivery of multiple agents and sustained antigen release, which may lead to synergistic interaction and stimulate durable effective cytotoxic T lymphocyte (CTL) responses against tumors^6^.

Improving the efficiency of cellular uptake remains a critical challenge for enhancing the therapeutic potential of NPs^7^. Inspired by pathogens^8,9^, researchers have designed virus-like nanomaterials with biomimetic topological structures to enhance the penetration of the complex cellular membranes and the uptake of immunotherapeutic agents^10^. The morphology of nanoparticles has a significant impact on cell membrane binding and instability. Sara Malekkhaiat Häffner et al found that the spikes on the surface of virus-like mesoporous silica (VLP) particles can disrupt the lipid bilayer of cell membranes^11^ and thus the VLP naturally possess enhanced cellular uptake. For example, Gao et al showed VLP mimic viral structures to enhance antigen delivery and immune activation. Co-loaded with imiquimod (IMQ) and ovalbumin (OVA), VLPs can improve cellular uptake and activate CD8^+^ T cells, effectively inhibit B16-OVA tumor growth and advance nonvaccine strategies for cancer immunotherapy. VLPs have superior biocompatibility in both cellular and in vivo models, enhanced cellular uptake, and a unique endocytic pathway that facilitated lysosomal escape and promoted antigen cross-presentation^12,13^. Though several types of Nano vaccines have been investigated in cancer immunotherapy, the mechanism of the VLPs in activating immune responses has remained poorly understood.

Proteomics is a powerful tool for the large-scale study of proteins to understand biological processes and disease therapy mechanisms^14^. It has been widely used in nanomedicine to identify protein corona formed on NPs and protein biomarkers for targeted drug delivery^15,16^. Since changes in protein expression are a key characteristic of immunotherapy, proteomics is also utilized to examine the immune interactions between cells and NPs. For example, Véronique Collin-Faure showed that the effects of polystyrene particles of varying sizes on macrophage polarization and inflammatory responses, the key proteins such as leupaxin or Pycard, under the innate immunity pathway from proteomic data were analyzed to explore the impact on immune responses^17^. The present study aimed to verify the activation of immune response with VLP, and further to discover the mechanism with proteomics analysis. The VLPs were prepared via microemulsion growth method^12^ and then functionalized with Polyethylene glycol (PEG) and amine silane. The VLPs were characterized by transmission electron microscopy (TEM), thermogravimetric analysis (TGA), and zeta potential measurements. The cytotoxicity, internalization efficiency, and immune response activation of the VLP were evaluated using *in vitro* cell viability assays, fluorescence microscopy, and flow cytometry, respectively. Finally, the immune response mechanism was studied with proteomics. Pathway enrichment was conducted, including pathway heatmaps from immunological databases, to identify immune-related pathways. Protein interaction networks and spike-dependent protein expression heatmaps were further analyzed. The present study may promote the fundamental understanding of the immune response triggered by VLPs, which offers new perspectives and directions to develop more efficient cancer immunotherapy.

## Materials and methods

### Preparation of the virus-like silica nanoparticles

Hexadecyltrimethylammonium bromide (CTAB, Sigma) was dissolved in deionized water with the concentration of 1%. 0.1 M sodium hydroxide was added slowly into the solution. The solution was stirred in an oil bath at 60 °C. Then, the mixture of tetraethyl orthosilicate (TEOS, Merck) and cyclohexane (Merck) with volume ratio=1:4 was added. The length of spikes on the VLPs were controlled by adjusting reaction time. The reaction mixture was stirred for 20, 48, and 70 hours to prepare the VLPs with the spike length of 0, 5 and 30 nm respectively. After the reaction, the VLPs were washed twice with ethanol to remove unreacted chemicals. And then, the VLPs were refluxed in a mixture of 150 ml ethanol at 99 °C for 48 hours to remove the surfactant template. The particles were stored in ethanol at an ambient temperature for further use.

For PEGylation, 2-[Methoxy(polyethyleneoxy)propyl] trimethoxy silane (90%, 6-9 PEG-units, abcr) was mixed with VLPs in ethanol and heated to 90 °C in an oil bath. After stirring for 30 minutes, the VLPs were washed twice with ethanol. 3-Aminopropyltriethoxysilane (APTES, VWR) was mixed with the PEG-modified VLPs and the resulting mixture stirred at 65 °C for 1 hour and finally washed with ethanol to remove unreacted chemicals. The samples were named PEG-VLP-NH_2_-0, PEG-VLP-NH_2_-5, and PEG-VLP-NH_2_-30, corresponding to the VLPs with the spike length of 0, 5, 30 nm, respectively.

### Characterization of VLPs

The particle morphology was characterized by using transmission electron microscopy (TEM). Diluted suspensions of VLPs were dropped onto carbon film grids (HC400-Cu, USA), dried in room temperature, and imaged using Jeol JEM-2100F TEM. The plain VLPs and VLPs after modification were measured by using Zetasizer Nano ZS (Malvern Panalytical Ltd, United Kingdom). Thermogravimetric analysis (TGA) was conducted to evaluate the amount of PEG functionalization on VLPs.

### Cell cytotoxicity of VLPs

RAW 264.7 cells were cultured at 37 °C with 5% CO_2_ in Dulbecco’s Modified Eagle Medium (DMEM, Biowest), supplemented with 10% fetal bovine serum (FBS, Sigma) and 1% penicillin-streptomycin (Gibco). When RAW 264.7 cells reached 80–90% confluence, they were washed with Hanks’ balanced salt solution (HBSS, Biowest) three times and detached with 0.25% Trypsin–EDTA (Gibco) for reseeding. For cell cytotoxicity measurements, the cells were seeded onto 96-well plates at a density of 1 × 10^4^ per well, cultured for 24 h, and incubated with the concentrations of VLPs (50 µg/mL) for 24 h. Untreated cells served as the positive control, and the cells treated with 10% Tween®20 as the negative control for cell viability. After incubation, the cells were washed three times with HBSS, 50 µL fresh cell culture medium was added into the wells, and then 50 µL ATP-based Cell Titer Glo assay (Promega) was added immediately. The plate was incubated at room temperature for 10 minutes, the luminescence was read by Fluoroskan Ascent FL microplate reader (Thermo Labsystems).

### Confocal microscopy of *in vitro* about cell uptake of VLPs

The RAW264.7 cells were seeded onto Ibidi 8-well plates with a cell density of 2 × 10^4^ cells per well and incubated for 24 h. The PEG-VLP-NH_2_ with varying spike lengths were labeled with Fluorescein 5(6)-isothiocyanate (FITC, Alfa Chemical) for internalization experiments. Specifically, 10 mg of VLPs after modification were stirred with 1 mg FITC in 2 mL of phosphate-buffered saline (PBS, VWR) for 12 hours at room temperature and covered from light. The FITC-labeled VLPs were collected by centrifugation and subsequently washed three times with PBS. And then the different spike lengths of FITC-labeled VLPs with 50 µg/mL in conditioned medium were added into wells and incubated for 4 h in the incubator. After the incubation, the cells were washed three times with HBSS to remove excess particles before staining with 5µg/mL CellMask™ red plasma membrane for 10 mins in the incubator. After the staining, the cells were washed twice with HBSS and fixed with 4% paraformaldehyde (Sigma-Aldrich) for 10 min at room temperature. For staining nucleus, the cells were stained with 10 µg/mL DAPI (4′,6-diamidino-2-phenylindole, Sigma-Aldrich) for 10 min at room temperature. Finally, the staining solution was replaced with HBSS. Fluorescence images were captured using a confocal laser scanning microscope instrument (Zeiss LSM 700).

### Quantitative proteomics

Raw 264.7 cells were seeded at 6×10^4^ cells per well onto six-well plates, incubated for 24 hours and then treated with PEG-VLP-NH_2_-0, PEG-VLP-NH_2_-5, and PEG-VLP-NH_2_-30 NPs (50 µg/mL) for 12 hours. To prepare Cell Extraction Buffer, Cell Extraction Buffer PTR (ab193970) and Cell Extraction Enhancer Solution (ab193971) were diluted with Milli-Q water and kept on ice. After washing the cells with cold PBS, 200 µL of Cell Extraction buffer was added per well and incubated on ice for 15 minutes. Lysates were centrifuged at 18 000 g for 20 min in a cold condition and supernatants (extracted solubilized proteins) were collected in separate Eppendorf low-protein binding tubes (Thermo Fisher Scientific). The protein content was quantified by the Pierce BCA Protein Assay Kit (Thermo Fisher Scientific, MA, USA) using Hidex microplate reader (Hidex, Finland).

The extracted and solubilized proteins were processed using SP3 protocol, as previously described^18^. Briefly, a total of 50 µg of solubilized proteins were reduced by dithiothreitol (DTT) (50 µg, Merk) for one hour while mixing at room temperature (RT), and alkylated by iodoacetamide (125 µg, Merk) for 30 minutes while mixing in the dark at RT. Reduced and alkylated proteins were then mixed with 100 µg of washed Sera-Mag magnetic carboxylate beads (Cytiva, 24152105050250) and 100 µg of washed Sera-Mag magnetic carboxylate beads (Cytiva, 44152105050250). An equal amount of pure ethanol (Sigma) was added to each tube and mixed at 1000 rpm and incubated for 10 minutes at RT. Beads were then pulled down on a magnetic rack for 2 minutes, prior to reconstitution in 140 µL of fresh 80 % ethanol and gently mixed. The magnetic separation and ethanol reconstitution was repeated 5 times before transferring the whole mixture to a new low protein binding Eppendorf tube in the last step. The protein samples were then digested on-beads using 1 µg of Pierce Trypsin Protease (Thermo Fischer) in 50 µL of ammonium bicarbonate buffer for 18 hours at 37º C while mixing at 1000 rpm in thermomixer (Thermo Fischer). The digested tryptic peptides were then recovered by magnetic separation and two additional washing steps with 50 µL of ammonium bicarbonate buffer. The recovered peptide solutions were then evaporated by SpeedVac vacuum concentrator (Thermo Fisher) at RT. The dried samples were reconstituted in a 2% acetonitrile acidified with 5 % formic acid, and mixed on a thermomixer at 300 rpm for 20 min in RT. Finally, 10 μl of each sample, containing 10 μg of digested proteins, was loaded for LC-MS/MS analysis in randomized order.

Proteomics analysis was performed using liquid chromatography tandem mass spectrometry (LC-MS) method as previously described^19^ using a Vanquish Flex UPLC system (Thermo Scientific) coupled to a high-resolution Orbitrap Q Exactive Classic mass spectrometer (Thermo Scientific) operating in positive ion mode. The mobile phase consisted of 0.1% formic acid (FA) in water (A) and 0.1% FA in acetonitrile (B). Peptides were loaded onto an Agilent AdvanceBio Peptide Map column (2.1 mm × 250 mm, 2.7 μm, Agilent Technologies) and separated at a flow rate of 0.3 mL/min using an 80-minute active gradient from 2% to 45% buffer B, with a total run time of 90 minutes. Mass spectrometric analysis was conducted using Full MS–SIM mode (resolution: 35,000; AGC target: 3 × 10^6^; maximum injection time: 60 ms; scan range: 385–1015 m/z) and data-independent acquisition (DIA) mode (resolution: 17,500; AGC target: 2 × 10^6^; maximum injection time: 60 ms; loop count: 25; isolation window: 24 m/z).

The proteomics raw data were analyzed using DIA-NN software (version 1.8.2) in library-free DIA mode^20^ with default parameters. The MS/MS spectral library and peptide retention times were predicted using the UniProt mouse reference proteome (UP000000589, updated February 2024, containing 21,957 protein entries, along with an additional Uniprot mouse database with 41,543 protein isoforms). Cysteine residues were treated as fixed modifications, while methionine oxidation and N-terminal acetylation were set as variable modifications, with a maximum of two variable modifications allowed per peptide. The predicted MS library was then used to search the raw data, applying a 1 % false detection rate threshold for both precursors and protein groups and requiring at least one proteotypic peptide of 7 to 30 amino acids in length. For data evaluation, the MaxLFQ normalized intensities^21^ were utilized. MaxLFQ intensities were processed and analyzed using Perseus software^22^. Protein group intensities were then log-transformed, and study groups were defined using the category-annotation tab. Data were categorized into two groups based on data completeness: a complete dataset (100% valid values) and a missing values dataset (at least three valid values in at least one group). For the complete dataset, categorical groups were compared using a linear model for microarray data analysis (LIMMA)^23^, followed by multiple testing correction using the false discovery rate (FDR), with statistical significance set at q-value < 0.05. For the missing values dataset, comparisons were conducted using a student’s t-test, considering statistical significance when at least three valid values were present in each group and a p-value < 0.01. Data visualization was carried out using R packages including dplyr, ggplot2, and heatmap. The Reactome database system was used to analyze gene-related metabolic pathways by Metascape^24^, and the pathway was analysis by SRPLOT^25^. In addition, STRING (Search Tool for the Retrieval of Interacting Genes/Proteins) was used to construct a protein-protein interaction network diagram, as described previously^26^.

### Flow cytometry of VLP

Human PBMC-derived monocyte-derived dendritic cells (MoDCs) were cultured in Roswell Park Memorial Institute (RPMI) medium supplemented with 10% fetal bovine serum (FBS) (Biowest), 1% penicillin-streptomycin (Gibco), and 2 mM L-Glutamine (Gibco). After incubating with the VLP for 16 hours, the collected cells were resuspended in 100 µL of flow cytometry buffer (1× PBS, 10% FBS [Gibco]) along with the appropriate concentrations of antibodies: HLADR-PE (BioLegend), CD86-PE-Cy7 (BioLegend), CD11c-FITC (BioLegend), CD40-APC (BioLegend), and CD80-BV510 (BD). The cells were incubated for 45 minutes at 4°C in the dark room. After incubation, the samples were washed twice with flow cytometry buffer and resuspended in buffer for analysis. Dendritic cells were collected and analyzed using flow cytometry on a NovoCyte Quanteon system (Biomedicum Flow Cytometry Unit). Data analysis was performed using FlowJo V.10.8.1, and graphs were generated using GraphPad Prism 10.3.0.

### Statistics and reproducibility

Data are presented as mean ± standard deviation (SD). The significant differences between the two groups were compared using ordinary one-way ANOVA analysis of the means of three or more unmatched groups followed by Dunnett’s method was used for multiple comparisons. The p-value is set in figures and legends such as: *P < 0.05, **P < 0.01 and ***P < 0.001.

## Results and discussion

### Physicochemical characterizations

The synthesis of VLP with different spike lengths was performed with the single-micelle epitaxial growth approach. The morphology of the VLPs was observed with TEM. The VLP-0 had a spherical shape (Figure 1a) while VLP-5 and VLP-30 presented virus like structure with respective spike length of 5 and 30 nm (Figure 1b, 1c), respectively. The average zeta potential for unmodified VLP-0, VLP-5, and VLP-30 was -24.5, -24.2, and -25.4 mV respectively. After polyethylene glycol (PEG) modification, zeta potential increased to -21.0 mV, -22.0 mV, and -20.5 mV for the PEGylated VLP with 0 nm, 5 nm, and 30 nm spike length, respectively. According to the weight loss in the TGA measurements, the amount of PEG modified on the VLPs was 5.0%, 4.3%, 3.7%, respectively (Figure 1e). PEG is widely used for modifying the surface of nanomaterials due to its neutrality, significant steric repulsion, and high hydrophilicity. By providing steric hindrance, PEG prolongs the blood circulation half-life of nanoparticles, thereby enhancing their biocompatibility^27^. After NH_2_ modification, surface charge reversed to be positive at +32.6 mV, +29.5 mV and +31.2 mV in the PEG-VLP-NH_2_-0, PEG-VLP-NH_2_-5, and PEG-VLP-NH_2_-30 respectively (Figure 1d). The positively charged amine groups can enhance drug loading efficiency through electrostatic interactions with various negatively charged biomolecules^28^, making them an essential component in immunization vaccine carrier applications.

**Figure 1.**
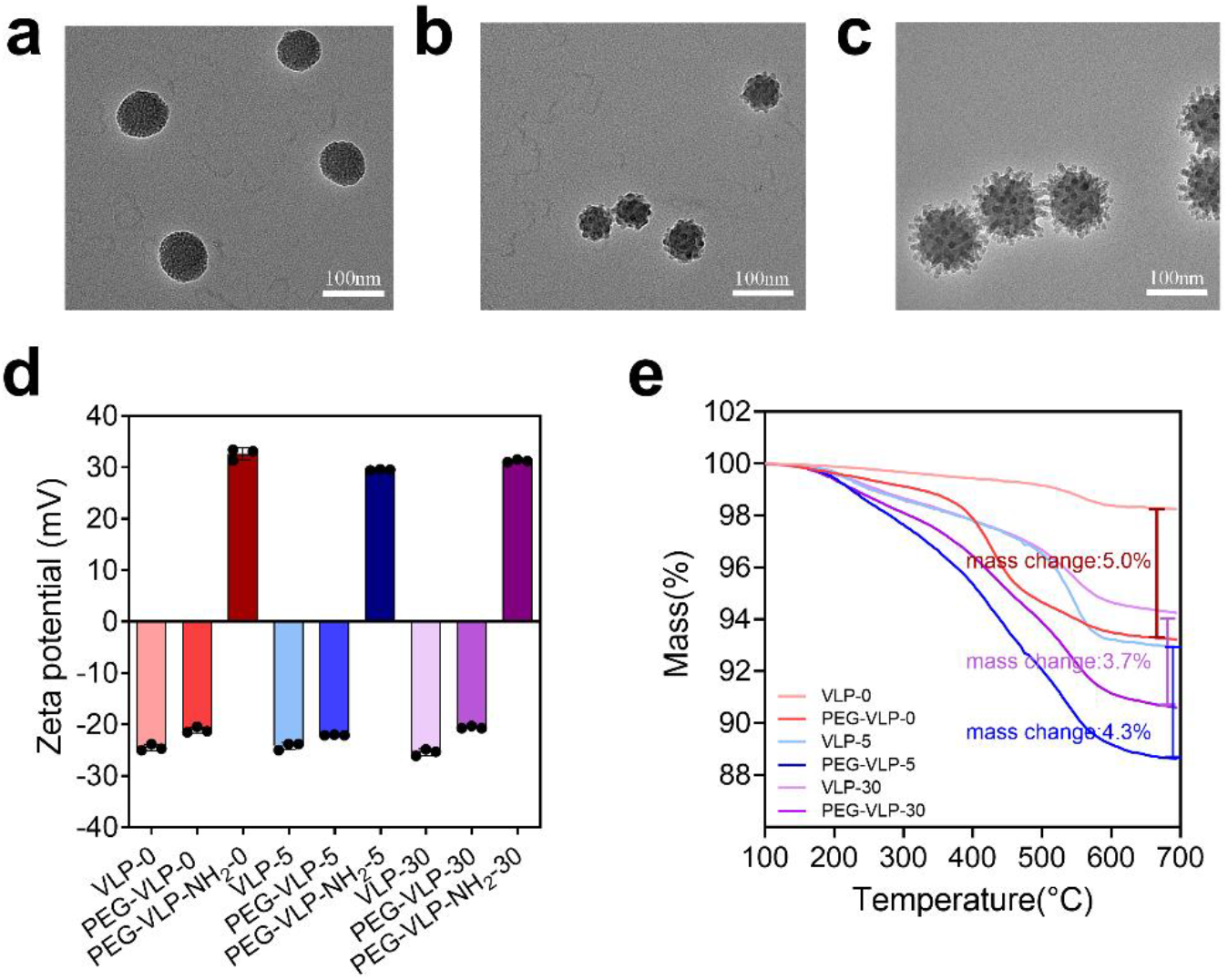
Characterizations of the VLP. a-c. TEM images of surface morphology of VLP with different spike lengths 0 nm, 5 nm, and 30 nm. The scale bar is 100 nm. d. Zeta potential of VLP before and after surface modifications. e. Thermogravimetric analysis (TGA) curves of VLPs to quantify PEG amount.

### Biocompatibility and cellular uptake of VLP

To investigate the biocompatibility of VLP cell viability of using RAW264.7 cells was measured after incubating the VLPs with the concentration of 50µg/mL. The results showed that the unmodified nanoparticles, VLP-0, did not cause significant cytotoxicity against RAW264.7. However, these unmodified VLP-5 and VLP-30 presented cytotoxicity at a concentration of 50 µg/mL reduced cell viability by 23.6% and 25.1%, respectively. After the VLPs were modified, PEGylated and amine-modified, the PEG-VLP-NH_2_-0 and PEG-VLP-NH_2_-5 did not have any impact on cell viability in the concentration of 50 µg/mL. The surface modified particles with the largest spike length, PEG-VLP-NH2-30, did not show cytotoxicity at 50 µg/mL. These findings show cytotoxicity with longer spikes but reduced cytotoxicity after PEGylation at the concentration of 50 µg/mL (Figure 2a).

**Figure 2.**
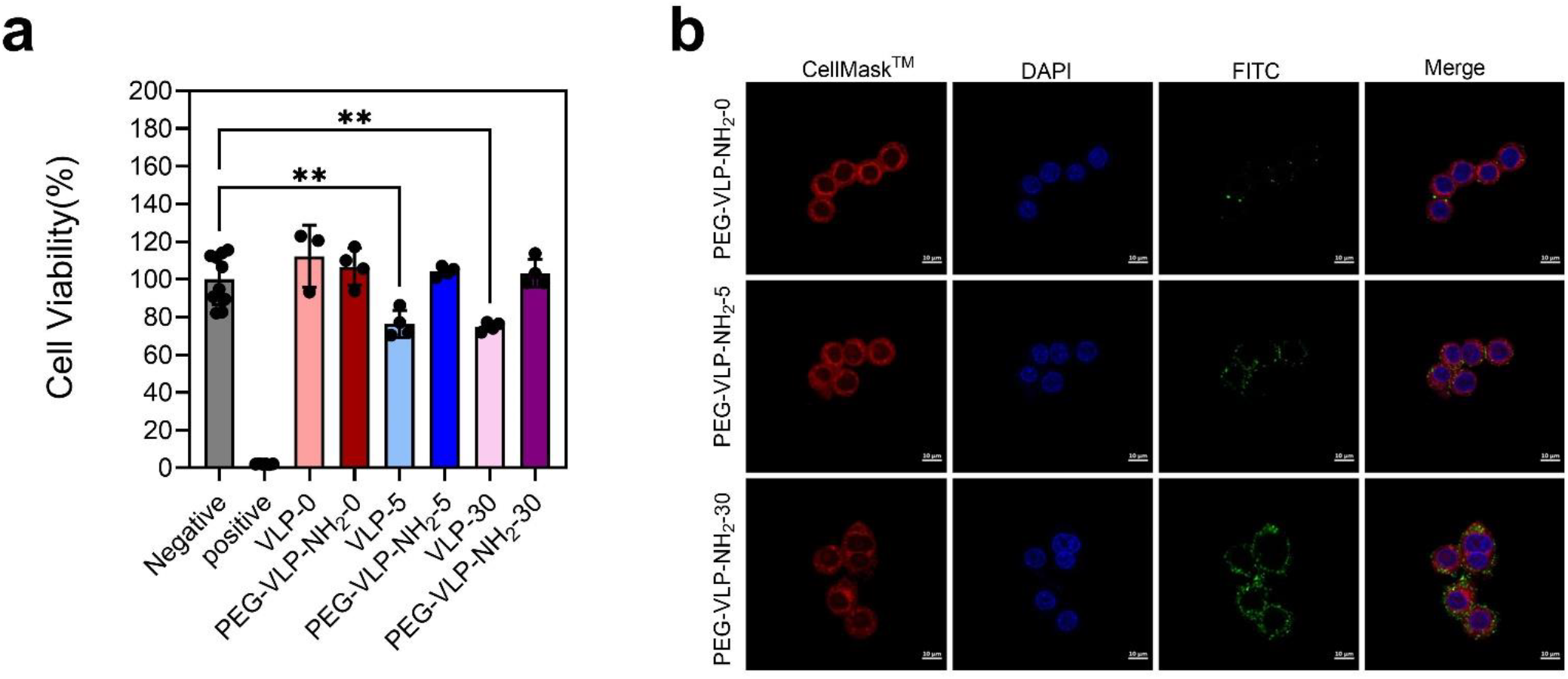
The cytotoxicity and uptake of virus-like mesoporous silica nanoparticles (VLP) into RAW264.7 cells at 50µg/mL. a. Viability of Raw264.7 cells treated with different VLPs after 24h at the concentration of 50µg/mL. Negative control=non-treated cells, positive control=lysed cells. The data represents mean ± SD. *, ** and *** represent statistically significant differences P < 0.05, P < 0.01 and P < 0.00, respectively. b. Fluorescence images of Raw264.7 cells treated with FITC-labeled VLPs of varying surface spike lengths. The VLP labeled with FITC (green color), the nucleus labeled with DAPI (blue color) and cell membrane labeled with CellMask^TM^ (red color). The scale bar is 10µm.

The cellular uptake efficiency of VLPs largely defines the efficacy of the immune response towards vaccines. Thus, the uptake of FITC-labeled VLPs was evaluated by fluorescence images with RAW264.7 cells. As shown in Figure 2b, the cellular uptake of VLPs appeared to correlate with the spike length, PEG-VLP-NH_2_-30 showing the most efficient, and PEG-VLP-NH_2_-0 the most modest uptake under the test conditions. Therefore, the result suggested that the enhancement in cellular uptake efficiency was mainly attributed to biomimetic spiky morphology, perhaps also explaining the association of cytotoxicity to the spike length.

### Protein expression of cells treated with VLP

String software served for studying Protein-Protein Interaction Networks to explore the relationships between differentially expressed proteins (Figure 3). With PEG-VLP-NH_2_-0, the upregulated protein network consisted of 71 nodes and 45 edges, with an average node degree of 1.27 and a local clustering coefficient of 0.44, forming 15 clusters. The downregulated proteins formed a smaller network with 37 nodes and 21 edges (average node degree: 1.14) and a local clustering coefficient of 0.39, clustering into 5 groups. Compared to PEG-VLP-NH_2_-0, PEG-VLP-NH_2_-5 exhibited an increase in both network size and connectivity. The upregulated proteins expanded to 79 nodes and 51 edges (average node degree: 1.29) while maintaining the same local clustering coefficient of 0.44, forming 16 clusters. The downregulated network showed an even more pronounced increase in connectivity, with 45 nodes and 52 edges (average node degree: 2.31), a local clustering coefficient of 0.435, and 6 clusters. The most significant expansion occurred in PEG-VLP-NH_2_-30, where the upregulated protein network reached 101 nodes and 62 edges, with an average node degree of 1.23 and a slightly lower local clustering coefficient of 0.43, forming 21 clusters. Notably, clusters related to immune pathways were also more visible in the sample of PEG-VLP-NH_2_-30. For example, the interaction cluster containing Traf1, Bcl10, and Tab1 which is associated with the TNFR1-induced NF-κB signaling pathway was found. The downregulated network also exhibited substantial growth, increasing to 65 nodes and 90 edges, with an average node degree of 2.77 and a higher local clustering coefficient of 0.49, forming 8 clusters. Overall, the results suggested the protein interaction networks varied across PEG-VLP-NH_2_-0, PEG-VLP-NH_2_-5, and PEG-VLP-NH_2_-30, showing progressive increases in node count, connectivity, and clustering complexity while increasing spike length.

**Figure 3.**
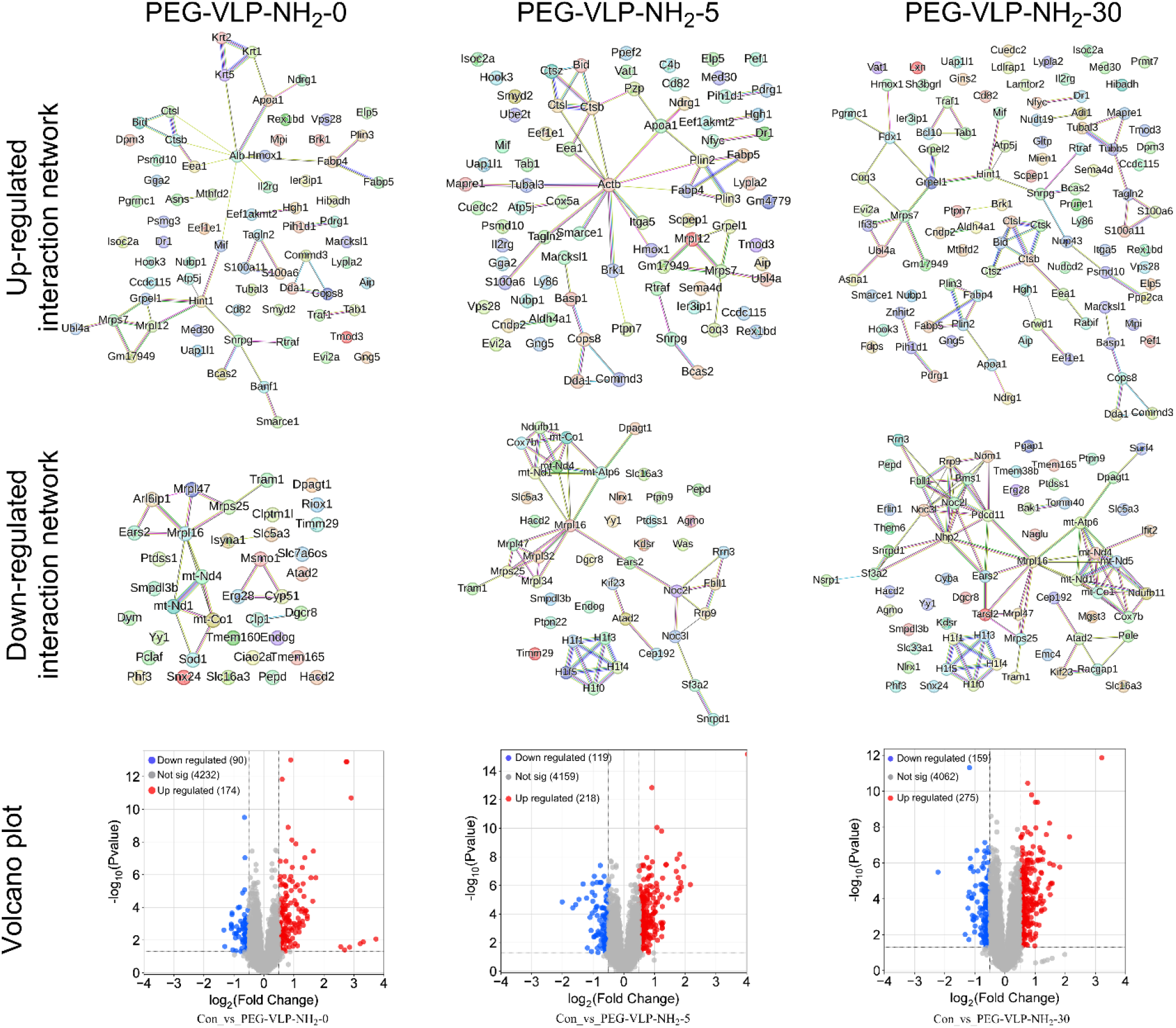
The differentially expressed proteins in RAW 264.7 were treated with PEG-NH_2_ modified VLPs as compared to untreated cell control. STRING Protein-Protein Interaction Networks of PEG-VLP-NH_2_-0, PEG-VLP-NH_2_-5, and PEG-VLP-NH_2_-30 with up and down regulated expressed protein. Volcano plots of expressed proteins from VLP with different spike lengths as compared to untreated cell control. The red dots represent the upregulated proteins, the blue dots represent the downregulated proteins, and the gray dots represent no significant change (P < 0.05, |log2FC|>0.5).

### Immune response related pathways induced by VLPs

Functional enrichment analysis was performed to study the pathways involved in the activation of immune cells by using Metascape with Reactome database. The pathways enriched at the highest level of statistical significance are shown in Figure 4 for each sample. PEG-VLP-NH_2_-0 treatment; upregulated keratinization and formation of the cornified envelope show high enrichment while expression of proteins involved in immune-related pathways were not shown. Treatment with PEG-VLP-NH_2_-5 induced upregulated proteins of pathways related to G alpha (s) signaling events are significantly enriched. Activation of the pathway stimulates adenylate cyclase to produce cyclic AMP (cAMP), consequently triggering various cellular signal transduction cascades ^29^.

**Figure 4.**
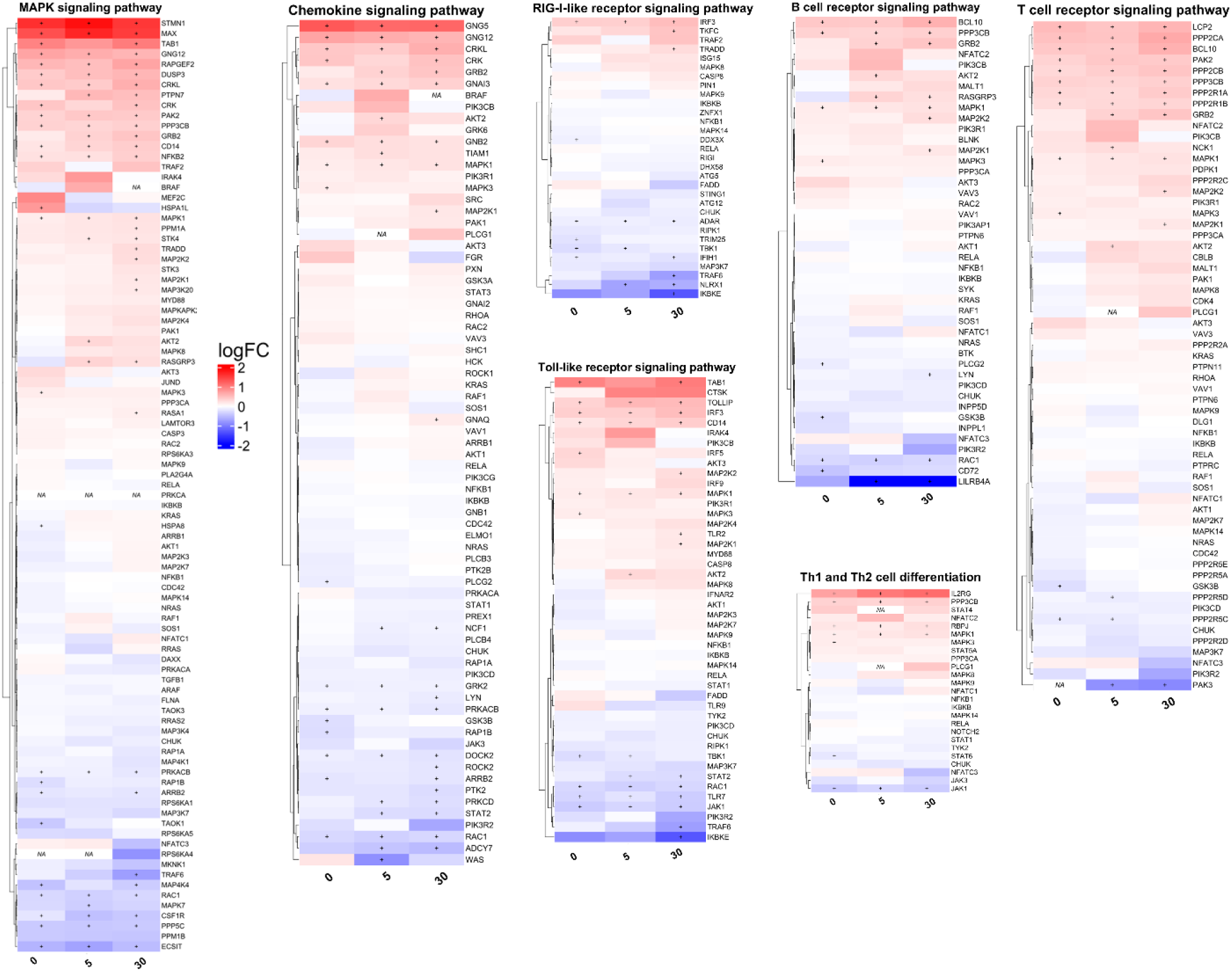
Pathway heatmap analysis after the cells incubated with three different spike length nanoparticles from an Immunological database. protein expressions are shown as logFC, + represents the p-value is lower than 0.01, red and blue colors correspond to significantly regulated proteins compared to the untreated control.

The treatment with PEG-VLP-NH_2_-30 induced upregulated proteins involved in trafficking and processing of endosomal Toll-like Receptor (TLR), TLR Cascades, and NF-kappa B signaling pathway are significantly enriched (Figure S2). TLRs activate a common signaling pathway, ultimately leading to the activation of the nuclear factor κB (NF-κB) transcription factor, as well as mitogen-activated protein kinases (MAPKs), including extracellular signal-regulated kinase (ERK), p38, and c-Jun N-terminal kinase (JNK). This cascade subsequently promotes the transcription of pro-inflammatory cytokines^30,31^. The down regulated protein for pathways enrichment analysis is shown in Figure S2 and does not include immune response related pathways. Heatmap of the results highlights the changes in expressed proteins of the Toll-like receptor signaling pathway quantified by logFC following treatments of cells with VLPs of varying spike lengths. The VLPs treatments altered the expressions patterns of proteins involved RIG-I-like receptors, chemokine, MAPK and TLR signaling pathways and Th1 and Th2 cell differentiation (Figure 4). Overall, the results imply the VLP spike length to stimulate the expression of various immune response related signaling pathways.

### The candidate immune related proteins induced by spike length

To study the effect of VLPs spike lengths for protein expression levels, we generated heatmaps of differences in protein expression levels between the treatment groups. Investigating the mechanisms underlying the increase in spike length is crucial, with a particular focus on upregulation proteins related to immune responses, such as TRAF6, and PIAS4 (Figure 5a) and downregulation of some proteins (Figure 5b). TRAF6 is a RING domain-containing E3 ubiquitin ligase and a member of the tumor necrosis factor receptor-associated factor (TRAF) family of adaptor proteins. It plays a key role in different signaling pathways and biological processes, serving as an important target for pro-inflammatory and immunoregulatory factors. Additionally, TRAF6 significantly enhances the transcriptional activation of nuclear factor κB (NF-κB), thereby playing an important role in the activation of innate immune responses^32^.TRAF6 is involved in multiple signaling pathways, and the Toll-like receptor 4 (TLR4) signaling pathway being particularly noteworthy. This pathway includes both MyD88-dependent and MyD88-independent mechanisms^33^, which subsequently cause a cascade of events leading to the activation of the MAPK pathway^34^, hereby inducing immune-related responses. This motivated us to look more closely at TRAF6 expressions because of VLP treatment. The results showed a statistically significant increase in the TRAF6 levels following treatment with PEG-VLP-NH_2_-30 (Figure 5c). Another protein we found of particular interest was PIAS4, a protein inhibitor of activated STAT4, plays a crucial role as a transcriptional coregulator in various cellular pathways. For example, PIAS4 may promote NF-κB activation by SUMOylating the inhibitor of NF-κB kinase subunit gamma (IKBKG)^35^, and then affect the immune response. The results showed PIAS4 upregulation to be statistically significant following PEG-VLP-NH2-30 treatment (Figure 5d).

**Figure 5.**
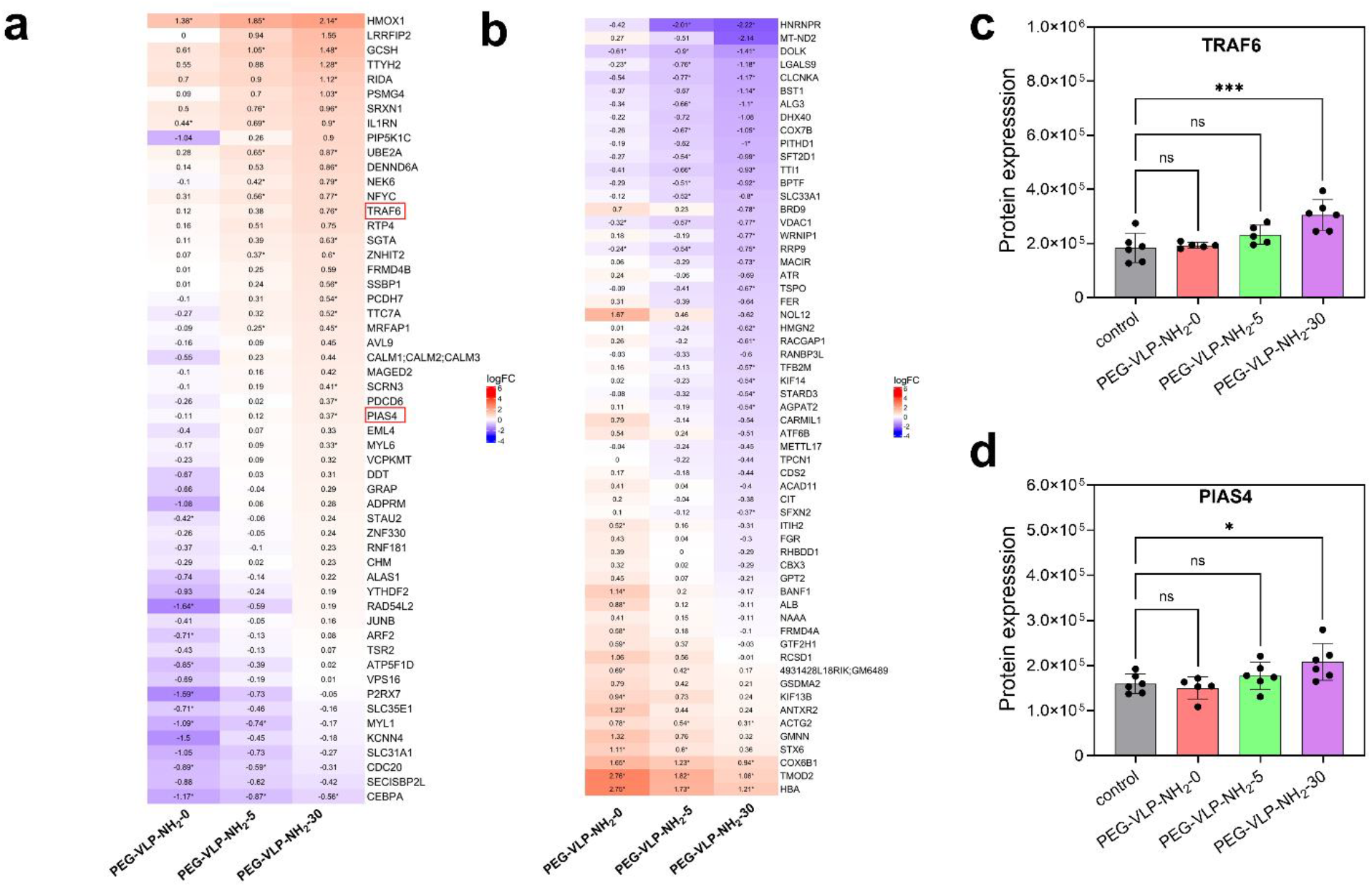
VLPs spike length-dependent protein regulation. a. Heatmap of proteins whose expression increased with the nanoparticle spiky length compared to the control. b. Heatmap of protein whose expression decreased with nanoparticle spiky length. Protein expressions are shown as logFC, red and blue colors correspond to significantly regulated proteins compared to the untreated control, the difference between each group is log2FC = 0.5. The significant protein expressions from proteomics analysis. c. TRAF6, d. PIAS4, *, ** and *** represent statistically significant differences P < 0.05, P < 0.01 and P < 0.001, respectively

### A**ctivation of immune response *in vitro***

Various surface markers on antigen-presenting cells, such as co-stimulatory receptors CD80, CD86, and CD40, dendritic cell marker CD11c, and a major histocompatibility complex (MHC class II) HLA-DR, are essential for T cell activation and immune regulation. The surface expression of these immune-related markers was studied with flow cytometry. The expressions of CD86 and CD80 were significantly upregulated with the PEG-VLP-NH_2_-30 compared to PEG-VLP-NH_2_-0 and PEG-VLP-NH_2_-5, indicating an excellent ability of VLP with long-spiky in activation of DCs (Figure 6a, 6b). CD80 and CD86 are co-stimulatory molecules for T cell activation and proliferation^36^. They are also markers for M1 macrophage activation^37^. The CD28/CD80 co-stimulatory signal activates pathways such as NF-kappa-B and MAPK, leading to the production of cytokines that initiate immune responses^38^. CD86 interacts with its ligands, CD28 or CTLA-4, to induce T lymphocyte proliferation and IL-2 production and is also involved in regulating B cell function^39,40^. HLA-DR, an MHC class II molecule, presents antigens to CD4^+^ T helper cells and is essential for initiating adaptive immune responses^41^. HLA-DR expression was significantly increased (p < 0.001) with PEG-VLP-NH_2_-5 and PEG-VLP-NH_2_-30 compared PEG-VLP-NH2-0 (Figure 6c). Furthermore, HLA-DR expression was upregulated with increased spike length (Figure 6d). The VLP did not induce a significant change in the expression of CD11c and CD40 regardless of spike length (Figure S3). These results suggested the VLPs have the potential to enhance antigen presentation to CD4^+^ T helper cells for cancer immunotherapy. Taken together, we conclude that VLPs effectively activate immune response, and the spike length plays a critical role in controlling the efficacy of immune stimulation.

**Figure 6.**
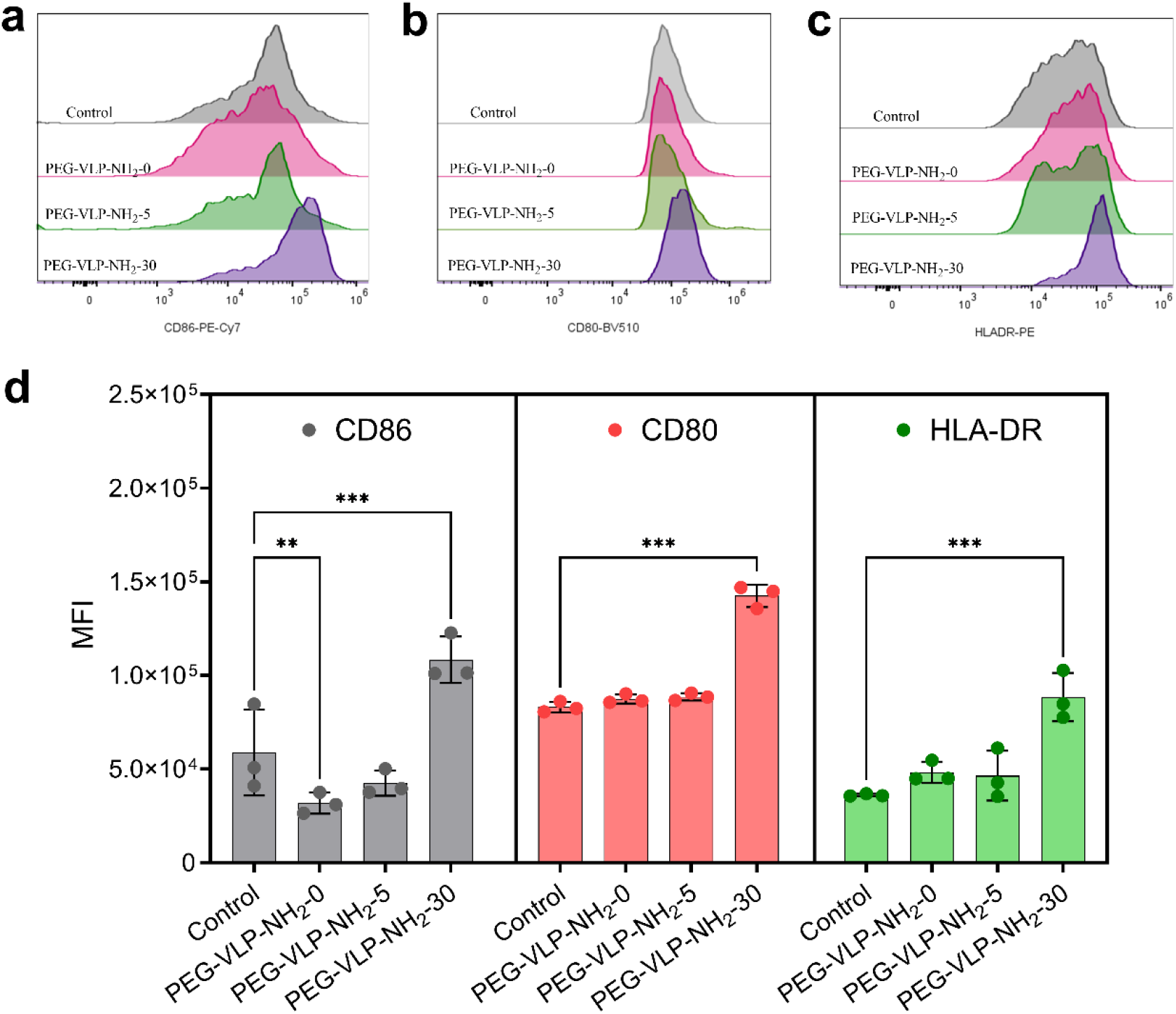
Flow cytometry analysis of CD80, CD86, and HLA-DR on human PBMC monocyte-derived dendritic cells (MoDCs) after being incubated with 50 µg/mL PEG-VLP-NH_2_-0, PEG-VLP-NH_2_-5, and PEG-VLP-NH_2_-30. a. CD86. b. CD80. c. HLA-DR. d. The violin plots quantify the expression levels. *, **, and *** indicate statistically significant differences with P < 0.05, P < 0.01, and P < 0.001, respectively.

## Conclusions

In this study, we investigated the effects of VLPs spike length on immune response in professional antigen-presenting cells. Our findings showed that particles with longer spikes (30 nm) showed higher cellular uptake efficiency than smooth mesoporous silica particles. Proteomic analysis indicated that VLPs with longer spikes more effectively induced immune response related proteins and pathways such as RIG-I-like receptors, chemokine, MAPK, Toll-like receptor signaling pathways, NF-κB signaling pathway and Th1and Th2 cell differentiation pathways. The expression levels of TRAF6 and PIAS4 proteins changed with spike length, further highlighting the impact of the spike-bearing VLPs on the immune responses. Furthermore, our results showed that the VLPs capability to upregulate the expression of CD86, CD80, and HLA-D, which is essential for stimulation of immune response by CD4^+^T cell. The finding of our study highlights biomimetic virus-like nanomaterials as potential immunoadjuvants for enhancing immunotherapy efficacy.

## Acknowledgments

This work was carried out with financial support from Research Council of Finland (Grant no. 356056, 359706, and 356992), program of China Scholarship Council, and Cancer Foundation Finland (Grant No. 230130). SIB Labs, UEF Cell and the Tissue Imaging Unit were acknowledged for technical support. We also acknowledge Biocentre Finland, Biocentre Kuopio for the use of their LC-MS laboratories.

## Declaration of Competing Interest

The authors declare that they have no known competing financial interests or personal relationships that could have appeared to influence the work reported in this article.

## Notes

### Competing Interest Statement

The authors have declared no competing interest.

### Summary of Updates

I deleted some data from the cell viability and kept the results about 50 ug/mL concentration.

## Reference

(1) Xu, B.; Li, S.; Shi, R.; Liu, H. Multifunctional Mesoporous Silica Nanoparticles for Biomedical Applications. Sig Transduct Target Ther 2023, 8 (1), 435. 10.1038/s41392-023-01654-7.

(2) Wen, H.; Tamarov, K.; Happonen, E.; Lehto, V.; Xu, W. Inorganic Nanomaterials for Photothermal-Based Cancer Theranostics. Advanced Therapeutics 2021, 4 (2), 2000207. 10.1002/adtp.202000207.

(3) Mody, K. T.; Popat, A.; Mahony, D.; Cavallaro, A. S.; Yu, C.; Mitter, N. Mesoporous Silica Nanoparticles as Antigen Carriers and Adjuvants for Vaccine Delivery. Nanoscale 2013, 5 (12), 5167. 10.1039/c3nr00357d.

(4) Yu, A.; Dai, X.; Wang, Z.; Chen, H.; Guo, B.; Huang, L. Recent Advances of Mesoporous Silica as a Platform for Cancer Immunotherapy. Biosensors 2022, 12 (2), 109. 10.3390/bios12020109.

(5) Wagner, J.; Gößl, D.; Ustyanovska, N.; Xiong, M.; Hauser, D.; Zhuzhgova, O.; Hočevar, S.; Taskoparan, B.; Poller, L.; Datz, S.; Engelke, H.; Daali, Y.; Bein, T.; Bourquin, C. Mesoporous Silica Nanoparticles as pHResponsive Carrier for the Immune-Activating Drug Resiquimod Enhance the Local Immune Response in Mice. ACS Nano 2021, 15 (3), 4450–4466. 10.1021/acsnano.0c08384.

(6) Escriche-Navarro, B.; Escudero, A.; Lucena-Sánchez, E.; Sancenón, F.; García-Fernández, A.; Martínez-Máñez, R. Mesoporous Silica Materials as an Emerging Tool for Cancer Immunotherapy. Advanced Science 2022, 9 (26), 2200756. 10.1002/advs.202200756.

(7) Zhang, R.; Qin, X.; Kong, F.; Chen, P.; Pan, G. Improving Cellular Uptake of Therapeutic Entities through Interaction with Components of Cell Membrane. Drug Delivery 2019, 26 (1), 328–342. 10.1080/10717544.2019.1582730.

(8) Sougrat, R.; Bartesaghi, A.; Lifson, J. D.; Bennett, A. E.; Bess, J. W.; Zabransky, D. J.; Subramaniam, S. Electron Tomography of the Contact between T Cells and SIV/HIV-1: Implications for Viral Entry. PLoS Pathog 2007, 3 (5), e63. 10.1371/journal.ppat.0030063.

(9) Chen, H.-J.; Hang, T.; Yang, C.; Liu, D.; Su, C.; Xiao, S.; Liu, C.; Lin, D.; Zhang, T.; Jin, Q.; Tao, J.; Wu, M. X.; Wang, J.; Xie, X. Functionalized Spiky Particles for Intracellular Biomolecular Delivery. ACS Cent. Sci. 2019, 5 (6), 960–969. 10.1021/acscentsci.8b00749.

(10) Xu, C.; Niu, Y.; Popat, A.; Jambhrunkar, S.; Karmakar, S.; Yu, C. Rod-like Mesoporous Silica Nanoparticles with Rough Surfaces for Enhanced Cellular Delivery. J. Mater. Chem. B 2014, 2 (3), 253–256. 10.1039/C3TB21431A.

(11) Häffner, S. M.; Parra-Ortiz, E.; Browning, K. L.; Jørgensen, E.; Skoda, M. W. A.; Montis, C.; Li, X.; Berti, D.; Zhao, D.; Malmsten, M. Membrane Interactions of Virus-like Mesoporous Silica Nanoparticles. ACS Nano 2021, 15 (4), 6787–6800. 10.1021/acsnano.0c10378.

(12) Wang, W.; Wang, P.; Tang, X.; Elzatahry, A. A.; Wang, S.; Al-Dahyan, D.; Zhao, M.; Yao, C.; Hung, C.-T.; Zhu, X.; Zhao, T.; Li, X.; Zhang, F.; Zhao, D. Facile Synthesis of Uniform Virus-like Mesoporous Silica Nanoparticles for Enhanced Cellular Internalization. ACS Cent. Sci. 2017, 3 (8), 839–846. 10.1021/acscentsci.7b00257.

(13) Gao, Y.; Zhang, Y.; Xia, H.; Ren, Y.; Zhang, H.; Huang, S.; Li, M.; Wang, Y.; Li, H.; Liu, H. Biomimetic Virus-like Mesoporous Silica Nanoparticles Improved Cellular Internalization for Co-Delivery of Antigen and Agonist to Enhance Tumor Immunotherapy. Drug Delivery 2023, 30 (1), 2183814. 10.1080/10717544.2023.2183814.

(14) Abdelkader, Y.; Perez-Davalos, L.; LeDuc, R.; Zahedi, R. P.; Labouta, H. I. Omics Approaches for the Assessment of Biological Responses to Nanoparticles. Advanced Drug Delivery Reviews 2023, 200, 114992. 10.1016/j.addr.2023.114992.

(15) Tenzer, S.; Docter, D.; Rosfa, S.; Wlodarski, A.; Kuharev, J.; Rekik, A.; Knauer, S. K.; Bantz, C.; Nawroth, T.; Bier, C.; Sirirattanapan, J.; Mann, W.; Treuel, L.; Zellner, R.; Maskos, M.; Schild, H.; Stauber, R. H. Nanoparticle Size Is a Critical Physicochemical Determinant of the Human Blood Plasma Corona: A Comprehensive Quantitative Proteomic Analysis. ACS Nano 2011, 5 (9), 7155–7167. 10.1021/nn201950e.

(16) Elechalawar, C. K.; Hossen, Md. N.; McNally, L.; Bhattacharya, R.; Mukherjee, P. Analysing the Nanoparticle-Protein Corona for Potential Molecular Target Identification. Journal of Controlled Release 2020, 322, 122–136. 10.1016/j.jconrel.2020.03.008.

(17) Collin-Faure, V.; Dalzon, B.; Devcic, J.; Diemer, H.; Cianférani, S.; Rabilloud, T. Does Size Matter? A Proteomics-Informed Comparison of the Effects of Polystyrene Beads of Different Sizes on Macrophages. Environ. Sci.: Nano 2022, 9 (8), 2827–2840. 10.1039/D2EN00214K.

(18) Hughes, C. S.; Foehr, S.; Garfield, D. A.; Furlong, E. E.; Steinmetz, L. M.; Krijgsveld, J. Ultrasensitive Proteome Analysis Using Paramagnetic Bead Technology. Molecular Systems Biology 2014, 10 (10), 757. 10.15252/msb.20145625.

(19) Montaser, A. B.; Gao, F.; Peters, D.; Vainionpää, K.; Zhibin, N.; Skowronska-Krawczyk, D.; Figeys, D.; Palczewski, K.; Leinonen, H. Retinal Proteome Profiling of Inherited Retinal Degeneration Across Three Different Mouse Models Suggests Common Drug Targets in Retinitis Pigmentosa. Molecular & Cellular Proteomics 2024, 23 (11), 100855. 10.1016/j.mcpro.2024.100855.

(20) Demichev, V.; Messner, C. B.; Vernardis, S. I.; Lilley, K. S.; Ralser, M. DIA-NN: Neural Networks and Interference Correction Enable Deep Proteome Coverage in High Throughput. Nature Methods 2020, 17 (1), 41–44. 10.1038/s41592-019-0638-x.

(21) Cox, J.; Hein, M. Y.; Luber, C. A.; Paron, I.; Nagaraj, N.; Mann, M. Accurate Proteome-Wide Label-Free Quantification by Delayed Normalization and Maximal Peptide Ratio Extraction, Termed MaxLFQ *. Molecular & Cellular Proteomics 2014, 13 (9), 2513–2526. 10.1074/mcp.M113.031591.

(22) Tyanova, S.; Cox, J. Perseus: A Bioinformatics Platform for Integrative Analysis of Proteomics Data in Cancer Research. In Cancer Systems Biology: Methods and Protocols; von Stechow, L., Ed.; Springer New York: New York, NY, 2018; pp 133–148. 10.1007/978-1-4939-7493-1_7.

(23) Ritchie, M. E.; Phipson, B.; Wu, D.; Hu, Y.; Law, C. W.; Shi, W.; Smyth, G. K. Limma Powers Differential Expression Analyses for RNA-Sequencing and Microarray Studies. Nucleic Acids Research 2015, 43 (7), e47–e47. 10.1093/nar/gkv007.

(24) Zhou, Y.; Zhou, B.; Pache, L.; Chang, M.; Khodabakhshi, A. H.; Tanaseichuk, O.; Benner, C.; Chanda, S. K. Metascape Provides a Biologist-Oriented Resource for the Analysis of Systems-Level Datasets. Nat Commun 2019, 10 (1), 1523. 10.1038/s41467-019-09234-6.

(25) Tang, D.; Chen, M.; Huang, X.; Zhang, G.; Zeng, L.; Zhang, G.; Wu, S.; Wang, Y. SRplot: A Free Online Platform for Data Visualization and Graphing. PLoS ONE 2023, 18 (11), e0294236. 10.1371/journal.pone.0294236.

(26) Hussein, H.; Kishen, A. Proteomic Profiling Reveals Engineered Chitosan Nanoparticles Mediated Cellular Crosstalk and Immunomodulation for Therapeutic Application in Apical Periodontitis. Bioactive Materials 2022, 11, 77–89. 10.1016/j.bioactmat.2021.09.032.

(27) Shi, L.; Zhang, J.; Zhao, M.; Tang, S.; Cheng, X.; Zhang, W.; Li, W.; Liu, X.; Peng, H.; Wang, Q. Effects of Polyethylene Glycol on the Surface of Nanoparticles for Targeted Drug Delivery. Nanoscale 2021, 13 (24), 10748–10764. 10.1039/D1NR02065J.

(28) Guo, Y.; Liu, M.; Cheng, Y.; Tian, J.; Feng, C.; Wang, Q.; Zhao, X.; Yin, L. Unraveling the Cytotoxicity and Cellular Pharmacokinetic of mPEG5-NH2 Polymers by UHPLC-MS/MS. Journal of Pharmaceutical and Biomedical Analysis 2025, 259, 116767. 10.1016/j.jpba.2025.116767.

(29) Kehrl, J. H. Heterotrimeric G Protein Signaling: Roles in Immune Function and Fine-Tuning by RGS Proteins. Immunity 1998, 8 (1), 1–10. 10.1016/S1074-7613(00)80453-7.

(30) Barton, G. M.; Medzhitov, R. Toll-Like Receptor Signaling Pathways. Science 2003, 300 (5625), 1524–1525. 10.1126/science.1085536.

(31) Kawai, T.; Ikegawa, M.; Ori, D.; Akira, S. Decoding Toll-like Receptors: Recent Insights and Perspectives in Innate Immunity. Immunity 2024, 57 (4), 649–673. 10.1016/j.immuni.2024.03.004.

(32) Bidère, N.; Snow, A. L.; Sakai, K.; Zheng, L.; Lenardo, M. J. Caspase-8 Regulation by Direct Interaction with TRAF6 in T Cell Receptor-Induced NF-κB Activation. Current Biology 2006, 16 (16), 1666–1671. 10.1016/j.cub.2006.06.062.

(33) Kawai, T.; Akira, S. The Role of Pattern-Recognition Receptors in Innate Immunity: Update on Toll-like Receptors. Nat Immunol 2010, 11 (5), 373–384. 10.1038/ni.1863.

(34) Li, J.; Liu, N.; Tang, L.; Yan, B.; Chen, X.; Zhang, J.; Peng, C. The Relationship between TRAF6 and Tumors. Cancer Cell Int 2020, 20 (1), 429. 10.1186/s12935-020-01517-z.

(35) Mabb, A. M.; Wuerzberger-Davis, S. M.; Miyamoto, S. PIASy Mediates NEMO Sumoylation and NF-κB Activation in Response to Genotoxic Stress. Nat Cell Biol 2006, 8 (9), 986–993. 10.1038/ncb1458.

(36) Sansom, D. M.; Manzotti, C. N.; Zheng, Y. What’s the Difference between CD80 and CD86? Trends in Immunology 2003, 24 (6), 313–318. 10.1016/S1471-4906(03)00111-X.

(37) Ishizuka, E. K.; Ferreira, M. J.; Grund, L. Z.; Coutinho, E. M. M.; Komegae, E. N.; Cassado, A. A.; Bortoluci, K. R.; Lopes-Ferreira, M.; Lima, C. Role of Interplay between IL-4 and IFN-γ in the in Regulating M1 Macrophage Polarization Induced by Nattectin. International Immunopharmacology 2012, 14 (4), 513–522. 10.1016/j.intimp.2012.08.009.

(38) Boulougouris, G.; McLeod, J. D.; Patel, Y. I.; Ellwood, C. N.; Walker, L. S. K.; Sansom, D. M. IL-2-Independent Activation and Proliferation in Human T Cells Induced by CD28. The Journal of Immunology 1999, 163 (4), 1809–1816. 10.4049/jimmunol.163.4.1809.

(39) Collins, A. V.; Brodie, D. W.; Gilbert, R. J. C.; Iaboni, A.; Manso-Sancho, R.; Walse, B.; Stuart, D. I.; Van Der Merwe, P. A.; Davis, S. J. The Interaction Properties of Costimulatory Molecules Revisited. Immunity 2002, 17 (2), 201–210. 10.1016/S1074-7613(02)00362-X.

(40) Lanier, L. L.; O’Fallon, S.; Somoza, C.; Phillips, J. H.; Linsley, P. S.; Okumura, K.; Ito, D.; Azuma, M. CD80 (B7) and CD86 (B70) Provide Similar Costimulatory Signals for T Cell Proliferation, Cytokine Production, and Generation of CTL. The Journal of Immunology 1995, 154 (1), 97–105. 10.4049/jimmunol.154.1.97.

(41) Gay, D.; Maddon, P.; Sekaly, R.; Talle, M. A.; Godfrey, M.; Long, E.; Goldstein, G.; Chess, L.; Axel, R.; Kappler, J.; Marrack, P. Functional Interaction between Human T-Cell Protein CD4 and the Major Histocompatibility Complex HLA-DR Antigen. Nature 1987, 328 (6131), 626–629. 10.1038/328626a0.

