## Appendix A. Supplementary information. for "Mechanism of biomimetic virus-like nanoparticles in triggering immune response elucidated by proteomics"

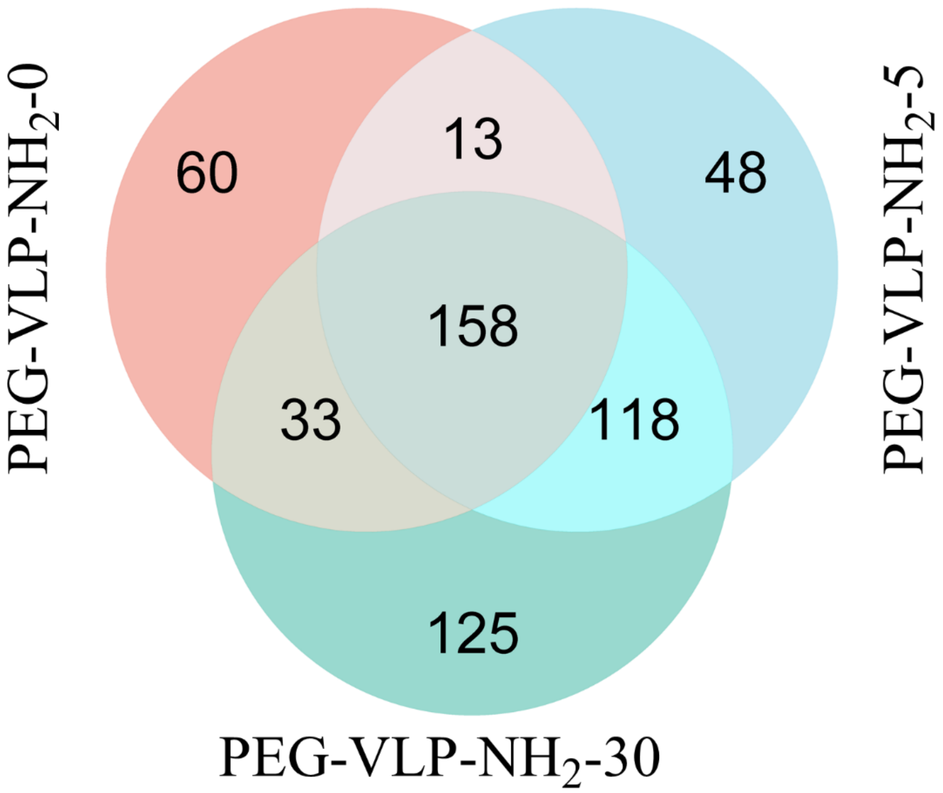


Figure S1. Venn diagram of differentially expressed proteins from VLP with spike lengths of 0, 5, and 30 nm compared to Control.

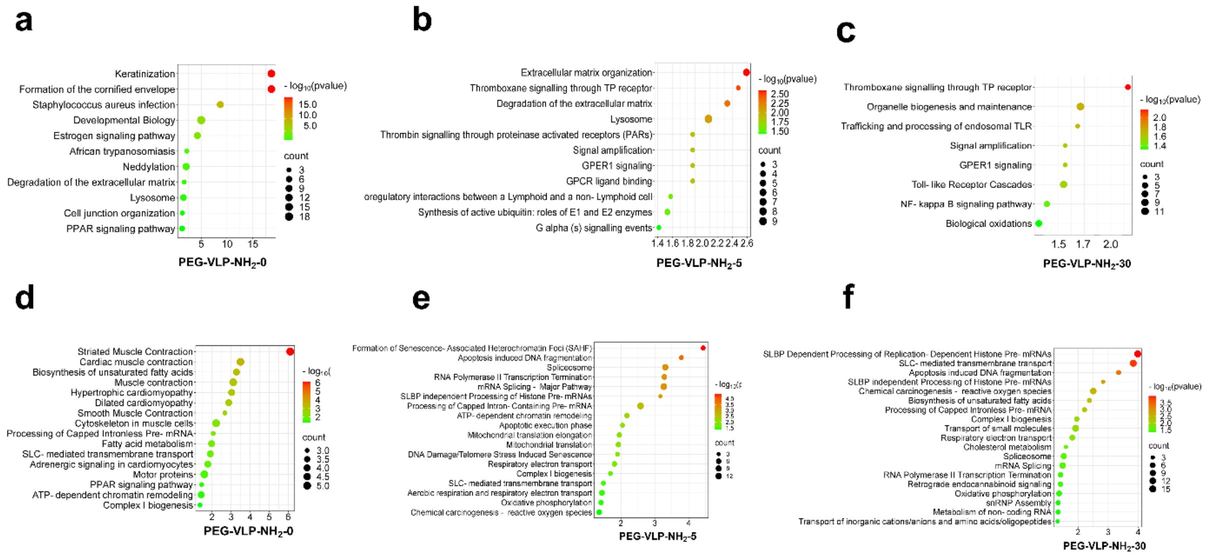


Figure S2. Pathway analysis of the cells treated with VLPs with varying spike length. a-c, Pathway enrichment analysis of up-regulated proteins from Reactome database. d-e, Pathway enrichment analysis of down-regulated proteins from Reactome database. Statistically significant enriched pathways are shown, the horizontal axis represents the gene ratio, while the vertical axis represents the enriched pathway name, the size of the dot reflects the count of proteins involved in that pathway, and the color scale indicates the statistical significance (-log10 (p-value)). d. Toll-like receptor signaling pathway heatmap from an Immunological database. + represents the p-value is lower than 0.01.

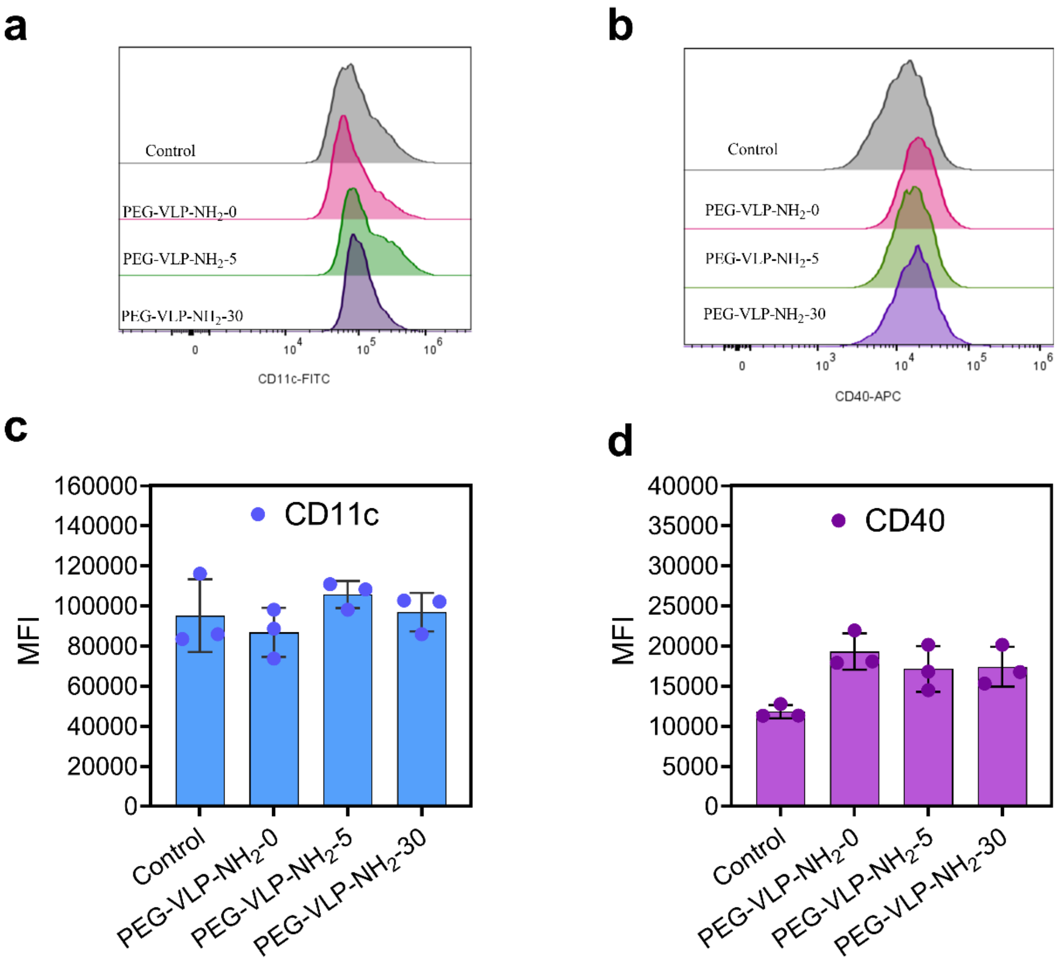


Figure S3. Flow cytometry analysis on human PBMC monocyte-derived dendritic cells (MoDCs) after being incubated with 50μg/mL PEG-VLP-$NH_2$-0, PEG-VLP-$NH_2$-5, and PEG-VLP-$NH_2$-30. a, c, CD11c. b, d, CD40.
